# NINJ1 regulates ferroptosis via xCT antiporter interaction and CoA modulation

**DOI:** 10.1101/2024.02.22.581432

**Authors:** Ssu-Yu Chen, Chao-Chieh Lin, Jianli Wu, Yubin Chen, Ya-En Wang, Yasaman Setayeshpour, Alexander Mestre, Jen-Tsan Chi

## Abstract

Ninjurin-1 (NINJ1), initially identified as a stress-induced protein in neurons, recently emerged as a key mediator of plasma membrane rupture during apoptosis, necrosis, and pyroptosis. However, its involvement in ferroptosis remains unknown. Here, we demonstrate that NINJ1 also plays a crucial role in ferroptosis, but through a distinct mechanism. NINJ1 knockdown significantly protected cancer cells against ferroptosis induced by xCT inhibitors but no other classes of ferroptosis-inducing compounds (FINs). Glycine, known to inhibit canonical NINJ1-mediated membrane rupture in other cell deaths, had no impact on ferroptosis. A compound screen revealed that NINJ1-mediated ferroptosis protection can be abolished by pantothenate kinase inhibitor (PANKi), buthionine sulfoximine (BSO), and diethylmaleate (DEM). These results suggest that this ferroptosis protection is mediated via Coenzyme A (CoA) and glutathione (GSH), both of which were found to be elevated upon NINJ1 knockdown. Furthermore, we discovered that NINJ1 interacts with the xCT antiporter, which is responsible for cystine uptake for the biosynthesis of CoA and GSH. The removal of NINJ1 increased xCT levels and stability, enhanced cystine uptake, and contributed to elevated CoA and GSH levels, collectively contributing to ferroptosis protection. These findings reveal that NINJ1 regulates ferroptosis via a non-canonical mechanism, distinct from other regulated cell deaths.

**Graphical abstract:** 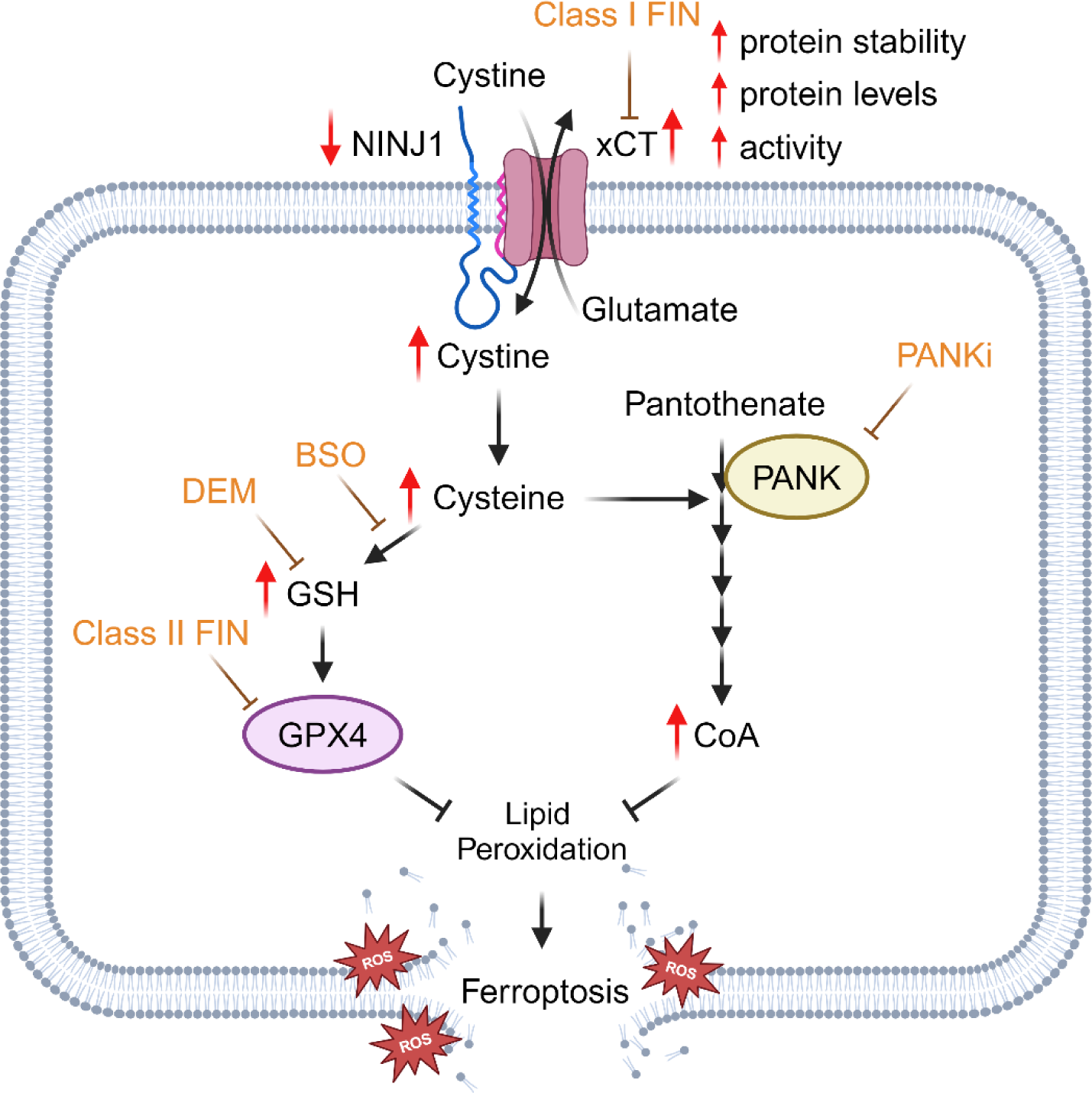

## Introduction

NINJ1 was initially discovered due to its induction upon nerve damage and found to promote neurite outgrowth and repair (1). Subsequent research has expanded our understanding of NINJ1’s functions, uncovering its role in cell adhesion (2), migration, and inflammation processes (3). Structurally, NINJ1 is a homophilic type II transmembrane protein conserved across species (4, 5). With extracellular and cytoplasmic domains, NINJ1 acts as a molecular bridge to transmit signals across the cell membrane to influence cell behavior in response to external stimuli. The role of NINJ1 in cancer is more complex, involving multiple processes that influence tumor progression, metastasis, and therapeutic response. Some studies show elevated NINJ1 levels in tumors and report a correlation with enhanced tumor progression and mobility (6, 7). Other research suggests that NINJ1 overexpression suppresses tumor growth (8–10) and its expression is downregulated during tumor recurrence and treatment resistance (11). This observation aligns with the involvement of NINJ1 in various cell death processes (4). Consequently, NINJ1 has been suggested as a biomarker and a therapeutic target in multiple human cancers.

Cell death plays a crucial role in both the biological growth and the pathophysiology of various diseases (12). Most regulated types of cell death involve protein pores disrupting osmotic balance, leading to water influx, cell swelling, and eventual cell membrane rupture. Different types of cell death involve different proteins, including the BCL family’s role in apoptosis (13), gasdermin D (GSDMD) in pyroptosis (14), and Mixed Lineage Kinase Domain-Like Protein (MLKL) in necroptosis (15), respectively. Interestingly, the NINJ1 protein was recently found to be involved in all three types of lytic cell death by aggregation to form a protein pore mediating the plasma membrane rupture (4, 16). The application of neutralizing antibodies targeting NINJ1 reduced cell death and prevented various tissue injuries (17). Additionally, glycine inhibits NINJ1 aggregation and blocks these types of cell death (18). These studies have firmly established NINJ1 as a critical and common executor of multiple cell death mechanisms by aggregating to form membrane-rupturing protein pores. However, the role of NINJ1 in ferroptosis was not discussed in these studies.

Ferroptosis is a regulated form of cell death characterized by iron dependency, lipid peroxidation, and oxidative stress (19, 20). Ferroptosis is suppressed by multiple endogenous antioxidant pathways including the System Xc-GSH-GPX4, trans-sulfuration, GCH1-BH4 pathway, DHODH-CoQH2 pathway, FSP1-CoQ10 pathway, and mevalonate pathway (21, 22). When these protective mechanisms are inhibited, iron-catalyzed oxidative damage of lipid membranes results in the rupture of the cellular membranes, leading to ferroptosis. Therefore, ferroptosis can be induced by diverse FINs, classified into distinct classes according to their targets and mechanisms. Class I (e.g., erastin, sulfasalazine (SAS)) targets xCT and cystine uptake (19, 23, 24), Class II (e.g., RSL3, ML162) targets GPX4 enzymatic activity (25), Class III (e.g., FIN56) depletes GPX4 and ubiquinone (26), and Class IV (e.g., FINO2) indirectly inhibits GPX4 while directly oxidizing iron (27–29). Ferroptosis has been appreciated to have significant implications for human diseases (21, 22). In cancer, ferroptosis presents an important tumor suppression mechanism. Since many treatment-resistant tumor cells have been found to be highly sensitive to ferroptosis (30, 31), the induction of ferroptosis has emerged as a promising therapeutic strategy (32).

While the involvement of NINJ1 in various types of cell death is well-established (4), its role in ferroptosis has remained elusive. A recent study demonstrated that NINJ1 is not essential in RSL3-induced ferroptosis (33). In this study, we systemically examined the role of NINJ1 in the ferroptosis of cancer cells induced by various FINs. While we were able to reproduce the finding that NINJ1 knockdown did not affect RSL3-induced ferroptosis, we found that NINJ1 knockdown robustly protected cancer cells against ferroptosis induced by xCT inhibitors, such as erastin and SAS. We conducted a compound screen to reveal that this protection against ferroptosis can be abolished by the chemical inhibition of the biosynthesis of either CoA (PANKi) or GSH (BSO and DEM), indicating the important role of both ferroptosis-protecting branches. Furthermore, we found that NINJ1 knockdown significantly elevated intracellular CoA levels and the GSH/GSSG ratio. CoA, similar to NINJ1 knockdown, protected ferroptosis triggered by xCT inhibitors, while showing no protection against GPX4 inhibitors. (24, 34). GSH, a well-known protector against ferroptosis, serves as a cofactor for GPX4. Furthermore, we found a physical association between NINJ1 and xCT. NINJ1 knockdown increased xCT stability, expression, and functionality that enhanced cystine uptake, resulting in an increase in the CoA and GSH levels. Conversely, NINJ1 overexpression specifically enhanced ferroptosis induced by xCT inhibitors, suggesting a therapeutic potential of NINJ1 in cancer through regulating ferroptosis.

## Results

### NINJ1 knockdown protected against ferroptosis specifically triggered by class I FINs

A previous study has shown that NINJ1 was not essential for RSL3-mediated ferroptosis (33). To systemically investigate the role of NINJ1 in ferroptosis, NINJ1 expression was successfully knocked down by several independent shRNA in HT-1080 human fibrosarcoma cells (Supplemental Fig. 1A-B). NINJ1-knockdown and control (empty vector) cells were then exposed to a dose-titration of various classes of FINs to compare their sensitivity to ferroptosis. We found that NINJ1 knockdown robustly protected HT-1080 cells against ferroptosis induced by class I FINs, such as erastin and SAS (Fig. 1A-B). In contrast, we observed that NINJ1 knockdown did not affect ferroptosis triggered by class II FIN (RSL3)-induced ferroptosis (Fig. 1C), which aligns with the recent study demonstrating the dispensability of NINJ1 in RSL3-induced ferroptosis (33). Similarly, NINJ1 knockdown did not affect ferroptosis induced by class III (FIN56) or class IV FINs (FINO2) (Fig. 1D-E). Collectively, these results indicate that NINJ1 regulates ferroptosis specifically induced by xCT inhibition mediated by class I FIN. Consistent with these phenotypic effects, NINJ1 knockdown also inhibited other erastin-induced molecular features of ferroptosis, including membrane rupture (Fig. 1F-G) and lipid peroxidation (Fig. 1H-I). A previous study showed that glycine protected cell death by inhibiting NINJ1 aggregation (18). Therefore, we also tested the effects of glycine and found it failed to confer protection against ferroptosis (Fig. 1J-K), suggesting that NINJ1 aggregation might not be involved in the protective effects against ferroptosis induced by NINJ1 knockdown. We also validated the ferroptosis-protective effects of NINJ1 knockdown in additional cancer cells representing different tumor types, including— MDA-MB-231 (breast cancer) and PC3 cells (prostate cancer) (Supplemental Fig. 1C-H). Collectively, these results suggest that NINJ1 knockdown specifically protects cancer cells against class I FINs, but not other classes of FINs.

**Figure 1.**
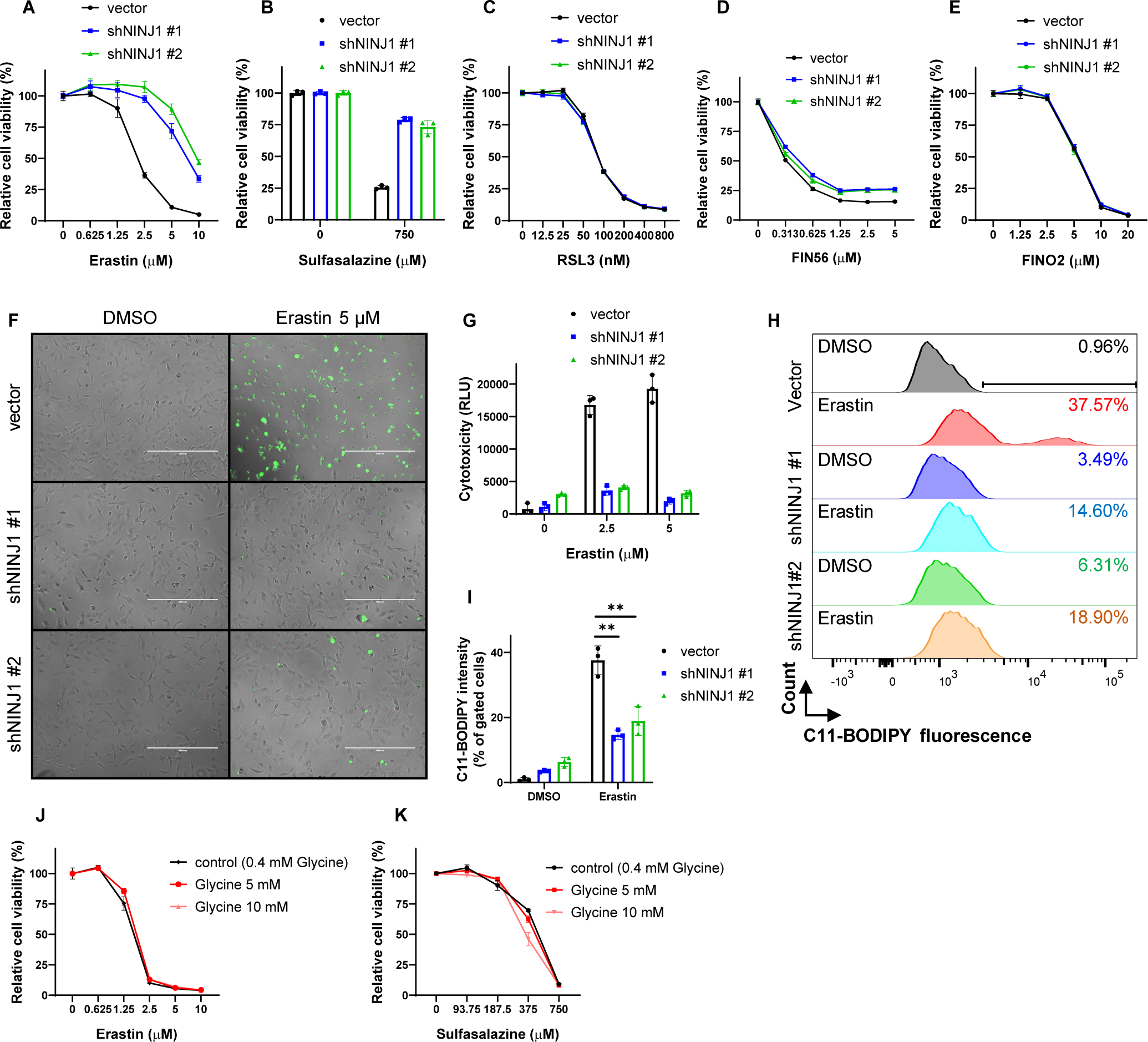
NINJ1 knockdown specifically protected against ferroptosis specifically triggered by class I FINs. (A-E) Cell viability of HT-1080 cells transduced with control vector and two NINJ1-targeting shRNA-mediated knockdown were determined by CellTiter-Glo assay after cells were treated with indicated concentrations of (A) Erastin (23 hours), (B) Sulfasalazine (28 hours), (C) RSL3 (19 hours), (D) FIN56 (19 hours), and (E) FINO2 (19 hours). (F-G) NINJ1 knockdown abolished Erastin-induced membrane rupture in HT-1080 cells. Membrane rupture of control and NINJ1-knockdown HT-1080 cells were observed by CellTox Green under fluorescence microscopy (F) and quantified (G) following Erastin treatment (2.5 and 5 μM, 22 hours). Scale bar: 400 μm. (H-I) NINJ1 knockdown inhibited erastin-induced lipid peroxidation in HT-1080 cells. Lipid peroxidation of vector and NINJ1-knockdown HT-1080 cells were determined by C11-BODIPY staining (H) and quantified by % of lipid peroxidation positive cells (I) following Erastin treatment (2.5 μM, 20 hours). (J-K) Cell viability of HT-1080 cells was determined by CellTiter-Glo assay after cells were treated with indicated concentrations of (J) Erastin (24 hours), (K) Sulfasalazine (48 hours) combined with Glycine (5 or 10mM).

### NINJ1 knockdown-mediated ferroptosis protection is abolished by the inhibition of CoA and GSH synthesis

The specificity of the NINJ1 knockdown protection phenotype against class I FINs suggested a mechanism upstream of the lipid repair enzyme, GPX4. To elucidate the potential mechanisms underlying this protection phenotype, we adopted a chemical genetic approach by conducting a focused compound screen that included compounds targeting ferroptosis-related pathways. Our goal was to identify compounds capable of abolishing this ferroptosis protection by reversing the relevant mechanisms. This involved treating both control and NINJ1 knockdown cells with DMSO or erastin, alone or in combination with various compounds. The relative cell viability, represented as a percentage of DMSO- or compound-treated cells under erastin treatment, is summarized as a heatmap (Fig. 2A), with detailed results for individual compounds presented in Fig. 2B-D and Supplemental Fig. 2. The examined compounds included etomoxir (35) (β-oxidation inhibitor, Supplemental Fig. 2A), lovastatin (26) (3-hydroxy-3-methylglutaryl-CoA reductase (HMGCR) inhibitor, Supplemental Fig. 2B), TOFA (36) (acetyl-CoA carboxylase (ACC) inhibitor, Supplemental Fig. 2C), dorsomorphin (36) (AMPK inhibitor, Supplemental Fig. 2D), alisertib (37, 38) (Aurora kinase A inhibitor, Supplemental Fig. 2E), verdinexor and leptomycin B (39) (XPO1/CRM1 inhibitors, Supplemental Fig. 2F-G), tipifarnib (40) (farnesyltransferase inhibitor, Supplemental Fig. 2H), methotrexate (41) (dihydrofolate reductase inhibitor, Supplemental Fig. 2I), pitstop2 (42) (clathrin-mediated endocytosis inhibitor, Supplemental Fig. 2J), EML425 and C646 (43) (p300/CBP Inhibitor, Supplemental Fig. 2K-L), brequinar (44) (dihydroorotate dehydrogenase (DHODH) inhibitors, Supplemental Fig. 2M), elamipretide (45) (mitochondrial-targeted peptide, Supplemental Fig. 2N), and the combination treatment of oligomycin and antimycin A (46) (mitophagy induction, Supplemental Fig. 2O). Remarkably, among all the tested compounds, only three compounds, including PANKi, BSO, and DEM, effectively abolished the observed ferroptosis protection mediated by NINJ1 knockdown (Fig 2A-D). The ferroptosis mitigation effects of these three compounds were further confirmed by all tested NINJ1-targeting shRNAs (Fig. 2B-D).

**Figure 2.**
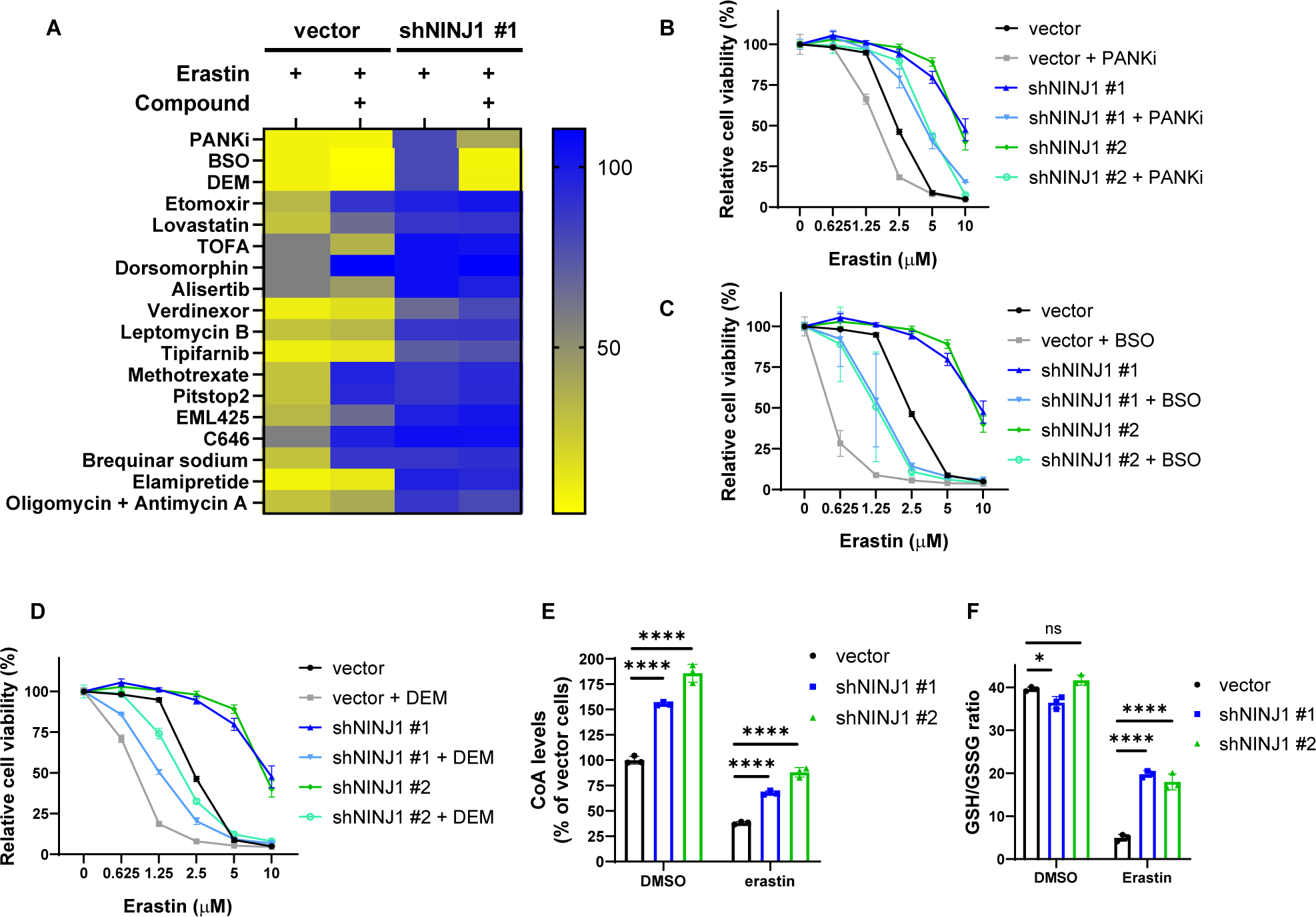
NINJ1 knockdown-mediated ferroptosis protection is abolished by the inhibition of CoA and GSH synthesis. (A) Heatmap showed the relative cell viability (% of DMSO or compound-treated cells) of control and NINJ1-knockdown HT-1080 cells were determined by CellTiter-Glo assay following treatment with Erastin combined with or without indicated compounds. (The cell viability graphs of individual compounds were listed in Fig. 2B-D and supplemental Fig. 2). (B-D) Cell viability of control and NINJ1-knockdown HT-1080 cells were determined by CellTiter-Glo assay after cells were treated with indicated doses of Erastin with or without (B) pantothenate kinase inhibitor (PANKi, 5 μM), (C) buthionine sulfoximine (BSO, 500 μM), and (D) diethylmaleate (DEM, 200 μM) for 22 hours. (B-D) were performed at the same time. Each group has the same control and NINJ1-knockdown HT-1080 cells following the treatment with indicated doses of Erastin. (E) Intracellular CoA levels of control and NINJ1-knockdown HT-1080 cells following Erastin treatment (1.25 μM, 23 hours) were determined by Coenzyme A Assay Kit. (F) GSH/GSSG ratio of contro and NINJ1-knockdown HT-1080 cells following Erastin treatment (2.5 μM, 19 hours) were determined by the GSH/GSSG-Glo Assay.

Pantothenate kinase (PANK) mediates the first and rate-limiting step of the CoA de novo biosynthesis pathway, which involves a five-step process utilizing cysteine (imported by xCT), pantothenate, and ATP (47). Recognizing its significance in this pathway, we evaluated intracellular CoA levels in HT-1080 cells following NINJ1 knockdown. Surprisingly, NINJ1 knockdown significantly elevated intracellular CoA levels before and after erastin treatment (Fig. 2E). Additionally, NINJ1 knockdown led to modest upregulation in the expression of PANK1 (Supplemental Fig. 2P-R), potentially contributing to increased CoA synthesis. Previous research has demonstrated the ability of CoA to rescue ferroptosis induced by xCT inhibition (24, 34). Results from our own studies, currently under review (48), also show that CoA addition rescues ferroptosis triggered by class I FIN (xCT inhibitors), but no other classes of FINs, consistent with the pattern of ferroptosis associated with NINJ1 knockdown.

The other two compounds which were found to mitigate ferroptosis protection were BSO and DEM. BSO is known for inhibiting γ-glutamylcysteine synthetase, a crucial enzyme mediating the initial step of GSH synthesis. Therefore, BSO is expected to impair GSH production (49). DEM is an electrophilic reagent that forms conjugates with GSH through the catalytic action of GSH transferase, leading to a rapid GSH depletion (50–52). Given these two compounds can abolish the protection phenotypes, we assessed GSH levels following NINJ1 knockdown. Although NINJ1 knockdown did not induce a significant alteration in the GSH/GSSG ratio without erastin, a noticeable increase in the GSH/GSSG ratio was observed in the NINJ1 knockdown group after erastin treatment (Fig. 2F). Taken together, these findings strongly indicate that both CoA and GSH elevation are essential for the NINJ1 knockdown-mediated ferroptosis protection phenotype.

### NINJ1 physically interacts with xCT

The synthesis of CoA involves the utilization of pantothenate, cysteine, and ATP (47), while GSH is synthesized from cysteine, glutamate, and glycine (53). Cysteine, a critical component in both CoA and GSH synthesis, is predominantly generated from the imported cystine (dimeric form of cysteine) into cells through the xCT transporter (54), which is the target of erastin. Since the ferroptosis inhibitory effect of NINJ1 knockdown was specific for xCT inhibitor, and considering that NINJ1 is also a membrane protein, we investigated its potential relationship with the xCT transporter. First, we determined their potential in vivo interaction at physiological levels using the proximity ligation assay (PLA), which revealed a significant co-localization signal when utilizing two antibodies recognizing NINJ1 and xCT (Fig. 3A). This interaction was specific, as the NINJ1 knockdown significantly reduced PLA signal and neither NINJ1 nor xCT antibody led to significant PLA signals (Fig. 3B-E). The physical association between NINJ1 and xCT was further confirmed using co-immunoprecipitation (Co-IP). Specifically, NINJ1-V5 and xCT-HA were expressed in HT-1080 cells. First, NINJ1 was pulled down using the V5 tag, separated on PAGE, transferred to a blot, and probed for the co-immunoprecipitated xCT, revealing a robust xCT signal (Fig. 3F). Reciprocally, when xCT was pulled down using the HA tag, separated on PAGE, transferred to a blot, and probed with NINJ1 to identify associated NINJ1 protein, a strong NINJ1 signal was detected (Fig. 3G), providing additional evidence of the physical association between NINJ1 and xCT. Furthermore, when EGFP-NINJ1 and mCherry-xCT were co-expressed in HT1080, fluorescent microscopy detected a merged yellow signal reflecting significant overlap between the green (NINJ1) and red (xCT) signals of these two membrane proteins (Supplemental Fig. 3A-C). Together, these findings strongly indicate a potential physical association between NINJ1 and xCT on the plasma membrane.

**Figure 3.**
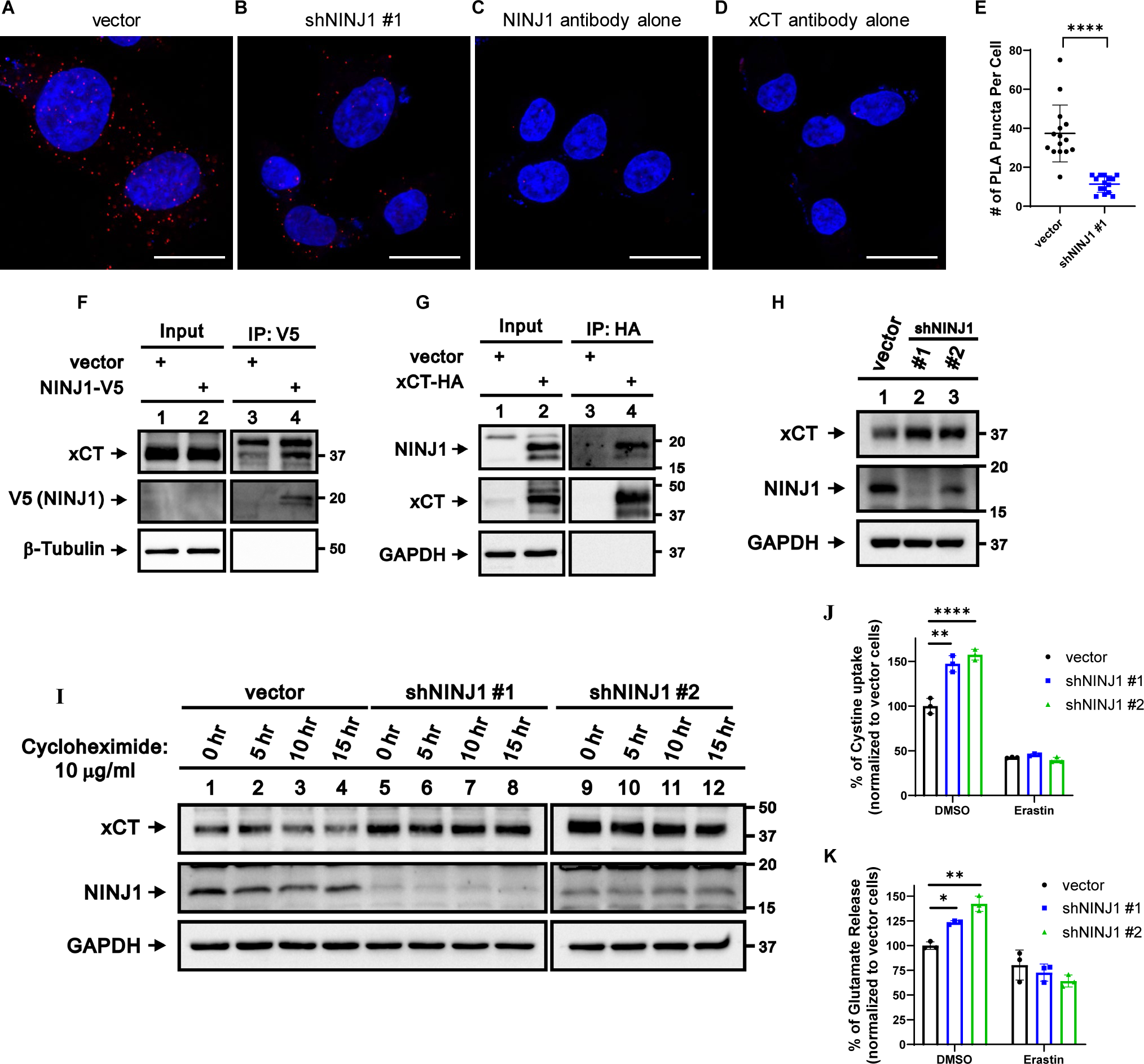
NINJ1 knockdown increased xCT stability, expression, and cystine import. (A-E) Control and NINJ1-knockdown HT-1080 cells were fixed in 3.7% paraformaldehyde for 15 min. PLA was subsequently conducted to examine the interaction between xCT and NINJ1. Representative images for the (A) vector, (B) NINJ1-knockdown, (C) NINJ1 antibody alone, and (D) xCT antibody alone conditions, along with (E) quantification of PLA signals, are presented. Scale bar: 20 μm. (F-G) Co-immunoprecipitation (CO-IP) and Western blot were used to detect the interaction between xCT and NINJ1. (F) V5-tag pulled down NINJ1 and (G) HA-tag pulled down xCT were probed using specified antibodies and subsequently detected through Western blot analysis. (H) xCT expression was increased following the NINJ1 knockdown in HT-1080 cells, which was verified by Western blot. (I) Enhanced xCT protein stability post-NINJ1 knockdown in HT-1080 cells was validated by cycloheximide treatment and subsequent Western blot analysis. (J) The levels of cystine uptake by the control and NINJ1-knockdown HT-1080 cells following Erastin treatment (10 μM, 4 hours) were determined by Cystine Uptake Assay Kit. (K) The levels of glutamate release by control and NINJ1-knockdown HT-1080 cells following Erastin treatment (1.25 μM, 20 hours) were determined by Amplex® Red Glutamic Acid/Glutamate Oxidase Assay Kit.

### NINJ1 knockdown increased the stability, expression, and activity of xCT

To determine the functional relevance of NINJ1 regarding xCT activity, we assessed the effects of NINJ1 knockdown on both the expression and functionality of xCT. NINJ1 knockdown in HT1080 led to a significant upregulation in xCT expression (Fig. 3H). Importantly, the enhanced xCT expression was consistently observed across various cancer cell lines, including MCF7, T47D, CAOV3, and PC3 cell lines (Supplemental Fig. 3D-G), highlighting the broad impact of NINJ1 knockdown. Additionally, we found that NINJ1 knockdown increased xCT stability, potentially contributing to elevated xCT protein levels (Fig. 3I).

Next, we wished to determine whether the increased xCT expression correlates with an enhanced xCT activity. Serving as an antiporter, xCT facilitates intracellular glutamate release and extracellular cystine uptake in a 1:1 ratio (54). We found that NINJ1 knockdown significantly increased cystine import (Fig. 3J) and glutamate export (Fig. 3K), indicating an enhanced xCT activity. Together, these data have that shown that NINJ1 knockdown increased the levels and function of xCT antiporter.

### NINJ1 overexpression specifically enhanced ferroptosis triggered by class I FINs

To further establish the connection between increased xCT expression/activity and the protective effects of NINJ1 knockdown, we overexpressed xCT (Supplemental Fig. 4A), measured cell viability following xCT inhibition, and assessed its activity. Consistent with the protective phenotype observed in NINJ1 knockdown, xCT overexpression conferred protection against erastin and SAS-induced ferroptosis (Fig. 4A-B), coupled with increased cystine import (Fig. 4C) and glutamate export (Fig. 4D). Collectively, these results suggest that the ferroptotic protective effects of NINJ1 knockdown may result from the enhanced xCT expression and activity, thereby increasing intracellular cystine levels and subsequent elevation of intracellular CoA and GSH.

**Figure 4.**
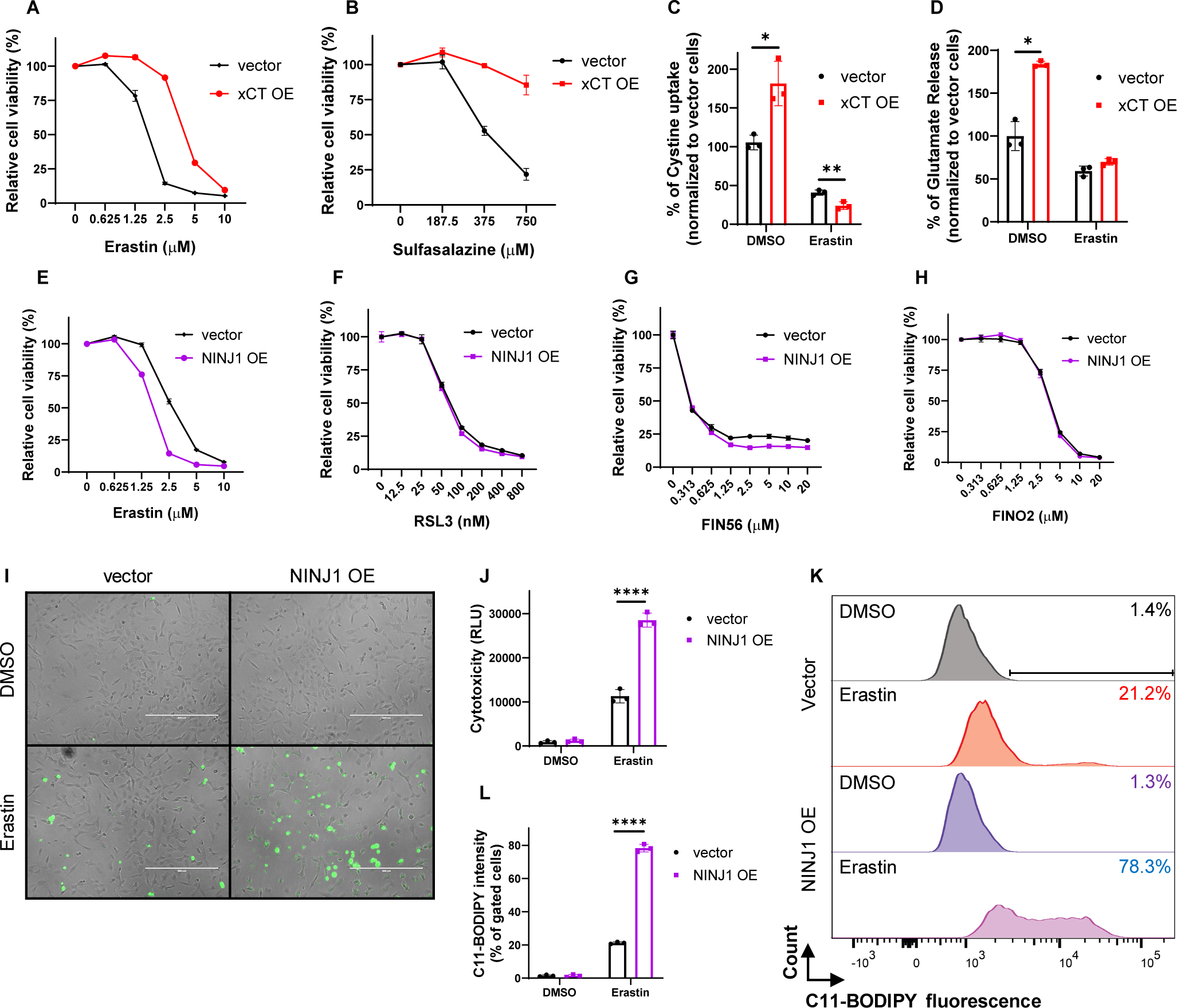
NINJ1 overexpression specifically enhanced ferroptosis triggered by class I FINs. (A-B) Cell viability of control and xCT-overexpression HT-1080 cells were determined by CellTiter-Glo assay after cells were treated with indicated concentrations of (A) Erastin (24 hours) and (B) Sulfasalazine (24 hours). (C) The levels of cystine uptake by the control and xCT-overexpression HT-1080 cells following Erastin treatment (10 μM, 4 hours) were determined by Cystine Uptake Assay Kit. (D) The levels of glutamate release by the control and xCT-overexpression HT-1080 cells following Erastin treatment (1.25 μM, 20 hours) were determined by Amplex® Red Glutamic Acid/Glutamate Oxidase Assay Kit. (E-H) Cell viability of control and NINJ1-overexpression HT-1080 cells were determined by CellTiter-Glo assay after cells were treated with indicated concentrations of (E) Erastin (23 hours), (F) RSL3 (17 hours), (G) FIN56 (17 hours), and (H) FINO2 (23 hours). (I-J) NINJ1 overexpression enhanced Erastin-induced membrane rupture in HT-1080 cells. Membrane rupture of control and NINJ1-overexpression HT-1080 cells were observed by CellTox Green under fluorescence microscopy (I) and quantified (J) following Erastin treatment (1.25 μM, 21 hours). Scale bar: 400 μm. (K-L) NINJ1 overexpression enhanced erastin-induced lipid peroxidation in HT-1080 cells. Lipid peroxidation of control and NINJ1-overexpression HT-1080 cells were determined by C11-BODIPY staining (K) and quantified by % of lipid peroxidation positive cells (L) following Erastin treatment (2.5 μM, 22 hours).

Next, we overexpressed NINJ1 in HT-1080 cells to assess its impact on ferroptosis (Supplemental Fig. 4B-C). In contrast to the outcomes observed with NINJ1 knockdown, our findings revealed that NINJ1 overexpression significantly enhanced ferroptosis induced by erastin (Fig. 4E), while exhibiting no significant impact on other classes of FINs (Fig. 4F-H). Corresponding to these observed phenotypic effects, NINJ1 overexpression also heightened other erastin-induced molecular features of ferroptosis, including membrane rupture (Fig. 4I-J) and lipid peroxidation (Fig. 4K-L). The NINJ1-mediated ferroptosis-enhancing effect was further confirmed in MDA-MB-231 cells (Supplemental Fig. 4D-G). Collectively, these results provide additional support for the ability of NINJ1 to regulate ferroptosis by class I FIN, but not other classes of FINs.

## Discussion

Based on these results, we proposed a model illustrating how NINJ1 regulates ferroptosis through a non-canonical mechanism (Fig. 5). Unlike conventional membrane pore formation by aggregated NINJ1 protein that disrupts the membrane, NINJ1 associates with xCT and regulates its level and function. Notably, NINJ1 knockdown enhances xCT level and activity by increasing xCT stability, resulting in elevated CoA and GSH levels, both of which were required to protect ferroptosis. Consequently, the disruption of either CoA or GSH synthesis abolishes the observed phenotypes of ferroptosis protection. Conversely, NINJ1 overexpression specifically intensified ferroptosis induced by xCT inhibitors, indicating a potential therapeutic role of NINJ1 that may enhance the response of cancer cells to ferroptosis.

**Figure 5.**
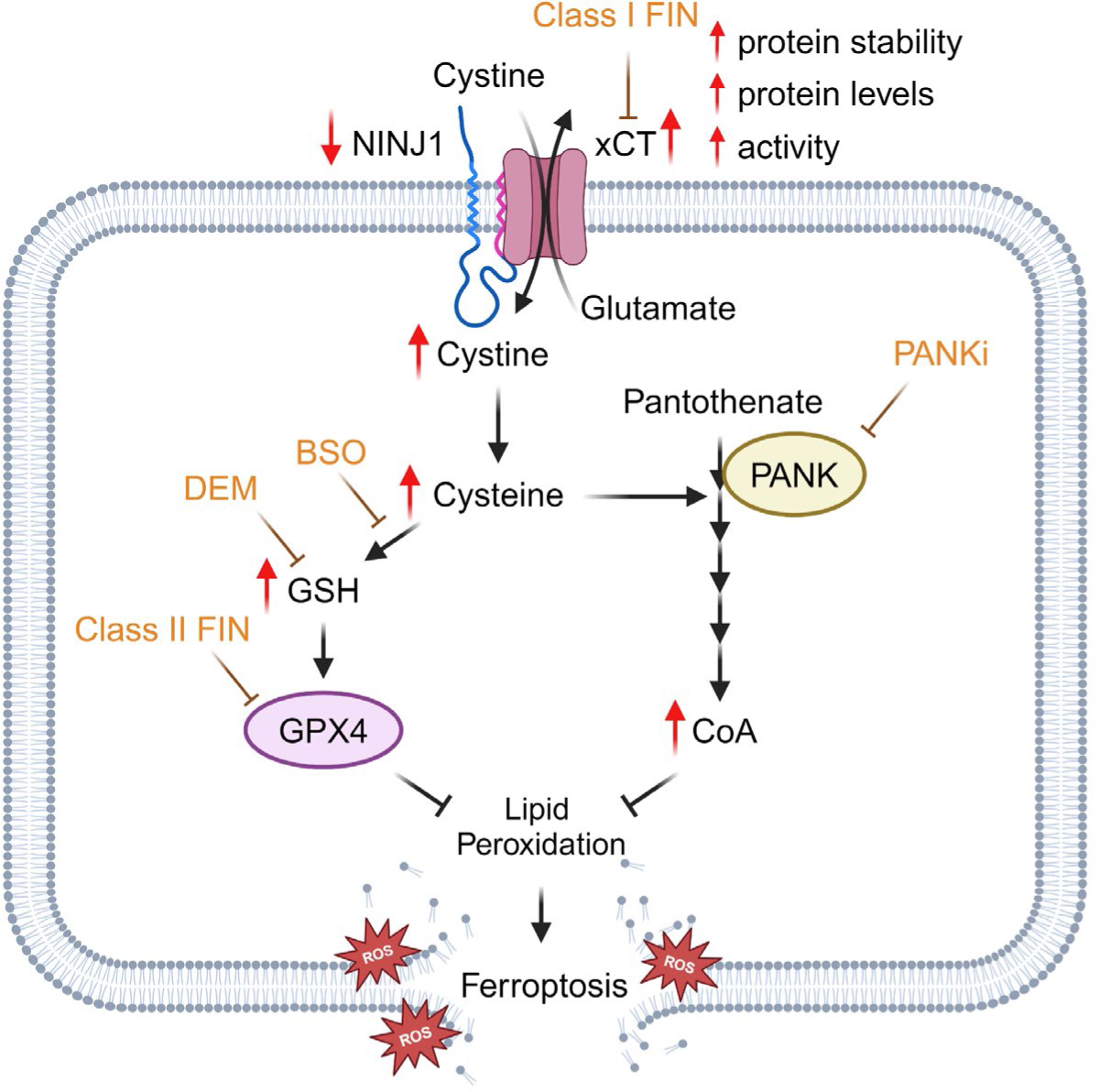
Proposed mechanism of the non-canonical role of NINJ1 in ferroptosis.

In our study, we identified a physical association between NINJ1 and xCT, and NINJ1 knockdown was found to increase the stability, expression, and functionality of xCT. These results suggest the potential role of NINJ1 in regulating the xCT degradation and restraining its function. Previous studies have shown that the protein stability of the xCT is regulated by various proteins, including solute carrier family 3 member 2 (SLC3A2), CD44 variant (CD44v), MUC1-C, EGFR, and OTUB1. Specifically, the system xc− consists of two integral subunits, xCT and SLC3A2, linked by a disulfide bond, wherein SLC3A2 functions as a chaperone responsible for maintaining xCT stability and ensuring its proper membrane localization (55). CD44v has been reported to interact with and stabilize xCT (56), and MUC1-C directly binds to CD44v, promoting xCT stability in the cell membrane (57). In glioma cells, the intracellular domain of EGFR has been reported to interact with and stabilize xCT (58). Additionally, OTUB1 also directly interacts with and stabilizes xCT, while the inhibitory effects of CD44 on ferroptosis are partly attributed to the enhancement of the OTUB1-xCT interaction (59, 60). Although we have shown that NINJ1 also regulates the stability of xCT, the underlying mechanisms and the potential involvement of other proteins in this regulation remain to be determined.

CoA and its thioester derivatives have been found to participate in various functions, such as the Krebs cycle, protein acetylation, amino acid metabolism, mitosis, fatty acid synthesis, and the regulation of gene expression (61, 62). In p53-mutant cells, CoA was initially recognized as a ferroptosis inhibitor, which showed that elevated CoA levels exhibited protective effects against erastin and glutamate-induced ferroptosis (34). In pancreatic tumor, CoA has been reported to enhance coenzyme Q_10_ production, contributing to ferroptosis protection (24). In our study, we observed that NINJ1 knockdown led to increased intracellular CoA levels, and the reversal of ferroptosis protection by PANKi validated the role of CoA in the ferroptosis protective effects initiated by NINJ1 knockdown. Previously reported data, along with our recent study in another manuscript, also suggest that CoA acts as a ferroptosis protective agent specifically against class I FINs in various cancer cell lines but does not exhibit the same protective effect against other classes of FINs. (24, 34, 48). This pattern of protection is consistent with the ferroptosis protection associated with NINJ1 knockdown. The increased CoA levels observed in NINJ1 knockdown may results from increased xCT functionality and cystine import, as well as increased PANK1 expression that may increase the de novo synthesis of CoA. Therefore, our results add to the increasing evidence that CoA is an important anti-ferroptosis metabolites subject to various regulators of ferroptosis.

While we analyzed ferroptosis in various cancer cell lines commonly used in ferroptosis research treated with different classes of FINs, it is possible that NINJ1 may play different roles in ferroptosis of other cell types and under other ferroptosis-triggering stimuli, such as nutrient deprivation, oxidative stresses, and ischemia reperfusion (63, 64). While our current research primarily focuses on the involvement of CoA and GSH, it is also plausible that additional mechanisms and metabolites may also contribute to ferroptosis protection. Consequently, there is still much to be explored regarding the role of NINJ1 in the ferroptosis of different cell types and under various ferroptosis-inducing conditions and disease processes.

Ferroptosis has been appreciated to have significant implications for human diseases (21, 22). For example, ferroptosis mediates cell death during acute kidney injury, stroke, cardiomyopathy (65), and various neurodegenerative diseases. Therefore, our study on the role of NINJ1 in ferroptosis suggests that modulating NINJ1, a membrane-accessible protein, holds a significant therapeutic potential. For example, various NINJ1 neutralization antibodies have been reported (17, 66, 67), and their application in these ferroptosis-associated diseases could potentially mitigate ferroptosis by inhibiting NINJ1. In human cancer, numerous studies highlight ferroptosis’s crucial role in tumor suppression and suggest its therapeutic potential in triggering tumor elimination and restraining tumor growth. For example, rapidly proliferating cancer cells may be more sensitive to ferroptosis given their upregulated intracellular iron levels (31), abnormal lipid metabolism, and increased ROS production (68, 69). Moreover, several processes associated with treatment resistance and metastasis, including epithelial-mesenchymal transition (EMT) (31), ECM signaling (30), and activation of YAP/TAZ (70), can promote ferroptosis (23). Therefore, targeting ferroptosis may provide important therapeutic options for tumors resistant to current treatments. In this context, various approaches for activating NINJ1 may have the potential to enhance ferroptosis and sensitize tumor cells to ferroptosis-inducing therapies.

## Methods

### Cell Culture

The HT-1080, MDA-MB-231, PC3, MCF7, T47D, and CAOV3 cells employed in this study were cultured following standard conditions. Specifically, they were cultured in a 37°C humidified incubator with 5% CO_2_. The culture media used were Dulbecco’s Modified Eagle Medium (DMEM) (GIBCO-11995) for HT-1080, MDA-MB-231, MCF7, T47D, and CAOV3, and Ham’s F-12K (Kaighn’s) Medium (GIBCO-21127022) for PC3. Additionally, the media were supplemented with 10% heat-inactivated FBS (Sigma, # F0926) and 10,000 UI/ml streptomycin and penicillin (ThermoFisher, #15140122).

### Constructs and lentivirus viral infections

To induce NINJ1 knockdown, we purchased small interfering RNAs (siRNAs) targeting human NINJ1 RNA from Dharmacon (D-017671-01-04), and two distinct shRNAs targeting human NINJ1 RNA were acquired from Sigma (TRCN0000063769, TRCN0000289088). For the overexpression of both NINJ1 and xCT, the pLX304-NINJ1 cDNA with a V5 tag was sourced from DNASU (HsCD00436123 and HsCD00438752), and the pWPI-SLC7A11 cDNA with an HA tag was obtained from Addgene (#201643). By using Gateway cloning, the NINJ1 cDNA from pENTR223-NINJ1 (DNASU, HsCD00505254) and SLC7A11 cDNA from pENTR223-SLC7A11 (DNASU, HsCD00512940) were individually subcloned into pLVpuro-CMV-N-EGFP (Addgene, #122848) and pLVpuro-CMV-N-mCherry (Addgene, #123221) for lentiviral expression. Lentiviruses carrying specific constructs were generated by transfecting HEK-293T cells with a mixture of the lentiviral vector, pMD2.G, and psPAX2 at a ratio of 1:1:0.1 using TransIT-LT1 transfection reagent (Mirus). The resulting blend was then filtered through a cellulose acetate membrane (0.45 µm, VWR, #28145-481). Subsequently, 250 µl of lentivirus-containing media and polybrene (8 µg/ml) was applied to a 60-mm dish of target cells and subjected to puromycin, blasticidin, or G418 selection.

### Chemicals

erastin (Bio-techne, 5449); SAS (MedchemExpress, HY-14655); RSL3 (Cayman, 19288); FIN56 (Cayman, 25180); FINO2 (MedchemExpress, HY-129457); PANKi (Cayman, 31002); BSO (Sigma, B2515); DEM (Sigma, D97703); etomoxir sodium salt (Selleckchem, S8244); lovastatin (Selleckchem, S2061); TOFA (Selleckchem, S6690); dorsomorphin (Selleckchem, S7306); alisertib (Selleckchem, S1133); verdinexor (Cayman, 26171); leptomycin B (Cayman, 10004976); tipifarnib (MedchemExpress, HY-10502); methotrexate (Selleckchem, S1210); pitstop2 (Sigma, SML1169); EML425 (Selleckchem, S2977); C646 (Selleckchem, S7152); brequinar (Selleckchem, S3565); elamipretide (Selleckchem, S9803)

### Cell viability and cytotoxicity

The CellTiter-Glo luminescent cell viability assay (Promega) was used to measure cell viability according to the manufacturer’s instructions. After the treatment, 10 μl of CellTiter-Glo reagent was applied to each well (containing 100 µl of media), shaken, and incubated for 10 minutes. The resulting luminescent signal was then quantified utilizing a chemiluminescence plate reader. Cytotoxicity quantification was performed using the CellTox Green assay (Promega) and a fluorescence plate reader. The dye was added to the media at a 1:1000 dilution.

### Lipid peroxidation assay

Lipid peroxidation was measured using C11-BODIPY (ThermoFisher Scientific, D3861) staining based on the manufacturer’s protocols. In brief, at the end of treatments, cells were incubated with 10 µM C11-BODIPY dye for 1 hour. Following harvest, washing, and resuspension in PBS with 1% BSA, lipid peroxidation levels were evaluated and quantified through flow cytometry (FACSCanto TM II, BD Biosciences).

### Western blots

Cells were lysed in RIPA buffer with protease inhibitor (Roche, 04693159001), constant vortexed at 4 °C for 30 minutes, and centrifuged at 16,000 rpm for 10 minutes. Supernatants were transferred to new tubes, and protein concentration was determined using the Pierce BCA protein assay kit (ThermoFisher, #23225). The western blot procedure involved loading the protein onto 15% SDS-PAGE gels, transferring the protein to a PDVF membrane, blocking the PVDF membrane for 1 hour with 5% non-fat milk in 1x TBST, and incubating the membrane with primary antibodies at 4 °C overnight. The primary antibodies included NINJ1 (1:1000, Invitrogen, PA5-95755), GAPDH (1:1000, Cell Signaling, #97166), PANK1 (1:1000, Cell Signaling, #23887), xCT (1:1000, Cell Signaling, #12691), V5 tag (1:1000, Cell Signaling, #13202), and β-tubulin (1:1000, Cell Signaling, #2128). NINJ1 or xCT proteins were isolated from HT-1080 cells overexpressing NINJ1-V5 or xCT-HA using the V5-tagged Protein Purification Kit Ver.2 (MBL, 3317) or HA tagged Protein Purification Kit (MBL, 3320), respectively. For the purification of both NINJ1 and xCT, HT-1080 cells were lysed in NP-40 buffer with protease inhibitor, subjected to constant vortexing at 4 °C, followed by centrifugation. The resulting supernatant was then purified using the V5 or HA-tagged Protein Purification Kit.

### Quantitative real-time PCR

By utilizing the RNeasy Mini Kit (Qiagen), RNA extraction was based on the manufacturer’s protocol. cDNA reverse transcription was conducted utilizing SuperScript IV reverse transcriptase (Invitrogen) and random hexamer. The synthesized cDNA, Power SYBR Green PCR Mix (Applied Biosystems), and primers were mixed following the manufacturer’s protocols, and quantitative real-time PCR reactions were performed on a StepOnePlus Real-time PCR system (Applied Biosystems). Absolute gene quantifications were normalized to GAPDH levels. The provided data represents three independent experimental repetitions. The primer sequences used were as follows: Human NINJ1 primers - sense, 5’-TCA AGT ACG ACC TTA ACA ACC CG −3’, antisense, 5’-TGA AGA TGT TGA CTA CCA CGA TG −3’; Human PANK1 primers - sense, 5’-TGG AAC GCT GGT TAA ATT GGT −3’, antisense, 5’-CCC AGT TTT CCC ATA AGC AGT AT −3’; Human GAPDH primers - sense, 5’-GGA GCG AGA TCC CTC CAA AAT −3’, antisense, 5’-GGC TGT TGT CAT ACT TCT CAT GG - 3’.

### Intracellular CoA levels

Intracellular CoA levels were quantified using the Coenzyme A Assay Kit (Sigma, MAK034) in accordance with the manufacturer’s instructions. In brief, cells were seeded in a 96-well plate and treated with the indicated drugs. Following the treatment, cells were washed with PBS, and each well received 40 μl of Coenzyme A Assay Buffer, 10 μl of the Coenzyme A Substrate Mix, and 2 μl of Conversion Enzyme Mix. After a 30-minute incubation at 37 °C, 50 μl of the master reaction mix, comprised of 2 μl Acyl-CoA Developer and 2 μl Fluorescent Peroxidase Substrate probe in Coenzyme A Assay Buffer, was added to each well and incubated for an additional 30 minutes. Intracellular CoA levels were then quantified using a fluorescent plate reader.

### GSH/GSSG ratio

To assess the GSH level, the GSH/GSSG-Glo Assay (Promega) was performed following the manufacturer’s protocols. In brief, after the designated treatment period, cells in the 96-well plate were washed with PBS and treated with 50 μl of either total GSH lysis reagent for total GSH measurement or oxidized GSH lysis reagent for GSSG measurement. Following this, all samples were added with 50 μl of Luciferin Generation Reagent and incubated for 30 minutes. Subsequently, 100 μl of Luciferin Detection Reagent was applied to all samples, incubated for an additional 15 minutes, and luminescence was measured. The GSH/GSSG ratios were directly determined from luminescence measurements, expressed in relative light units (RLU).

### Proximity Ligation Assay

The interaction between NINJ1 and xCT proteins was assessed using the Duolink Proximity Ligation Assay (Sigma) following the manufacturer’s protocol. Initially, cells were fixed with 3.7% formaldehyde, permeabilized using 0.3% Triton X-100, and blocked with Duolink Blocking Solution for 1 hour. Primary antibodies targeting NINJ1 (1:200, Invitrogen, PA5-95755) and xCT (1:200, Invitrogen, MA5-44922) were applied to each sample and incubated overnight at 4 °C. Subsequent to washing with 1x Wash Buffer A, PLUS and MINUS PLA probes were introduced to each sample and incubated at 37 °C for 1 hour. Following this step, after a thorough wash, ligation solutions were administered and allowed to incubate at 37 °C for 30 minutes. After three additional wash cycles, amplification solutions were added and incubated at 37 °C for 100 minutes. Following two washes with 1x Wash Buffer B and one wash with 0.1x Wash Buffer B, the slides were mounted using Duolink PLA Mounting Medium with DAPI for nuclear staining and then covered with a coverslip. Immunofluorescence microscopy was performed with a confocal microscope featuring Airyscan for super-resolution quality (880, Zeiss).

### Cystine uptake assay

The xCT activity was evaluated using the Cystine Uptake Assay Kit (DOJINDO, UP05) following the manufacturer’s protocol. In summary, after completing the treatment, cells in a 96-well plate underwent three washes with cystine-free and serum-free medium, followed by incubation in cystine-free and serum-free medium at 37 °C for 5 minutes. Subsequently, cystine uptake solution (with DMSO or erastin) or cystine-free and serum-free medium (blank), was added to each sample and incubated at 37 °C for 30 minutes. After three washes with PBS, methanol and the working solution were added to each sample, thoroughly mixed, and then incubated for an additional 30 minutes at 37 °C. The fluorescence intensity of each sample was measured using a fluorescence plate reader. After normalizing the fluorescent signal to the total cell number at the end of the experiment, the levels of cystine uptake was expressed as a percentage of the control group with no treatment (DMSO).

### Glutamate release assay

The release of intracellular glutamate into the extracellular medium was quantified using the Amplex Red Glutamic Acid/Glutamate Oxidase Assay Kit (Invitrogen, A12221) following the manufacturer’s protocol. Specifically, upon completing the treatment, cells in 6-well plates were subjected to two washes with PBS and then incubated in Na+-containing and glutamine-free media containing either DMSO or erastin for 1 hour. Subsequently, 50 μl of the medium from each well was transferred to a 96-well plate and incubated with 50 μl of a reaction mixture for 30 minutes. The fluorescence intensity of each sample was then measured using a fluorescence plate reader. After normalizing the fluorescent signal to the total cell number at the end of the experiment, the amount of glutamate release was expressed as a percentage of the control group with no treatment (DMSO).

### Statistics

Bar graphs include individual data points to represent biological replicate numbers, while figure legends for the line graph specify the number of biological replicates. Data are expressed as the mean +/- standard error of the mean (SEM). Statistical analyses were performed using a two-tailed Student’s t-test or one-way ANOVA with Tukey’s multiple comparisons in IBM SPSS Statistics 21 version or Prism (Graphpad). Error bars indicate SEM, and significance between samples is indicated as follows: *p < 0.05, **p < 0.01, ***p < 0.001, and ****p < 0.0001.

### Data availability

All data and reagents supporting the findings of this study are available from the authors upon reasonable request.

## Author contributions

S.Y.C. and J.T.C. conceived the experiments and wrote the manuscript. S.Y.C. performed the majority of the experiments. J.T.C. supervised the work. C.C.L., J.W., Y.C., Y.E.W., Y.S., and A.M. collaborated in the discussion and experiments.

## Acknowledgments

We are grateful for the technical support from the members of the Chi lab. We acknowledge the financial support in part by DCI Pilot Project, DOD grants (OC230018, W81XWH-17-1-0143, W81XWH-15-1-0486, W81XWH-19-1-0842, W81XWH-20-1-0907) and NIH grants (R01GM124062, 1R21-AI149205).

