## Supplemental Figures for "NINJ1 regulates ferroptosis via xCT antiporter interaction and CoA modulation"

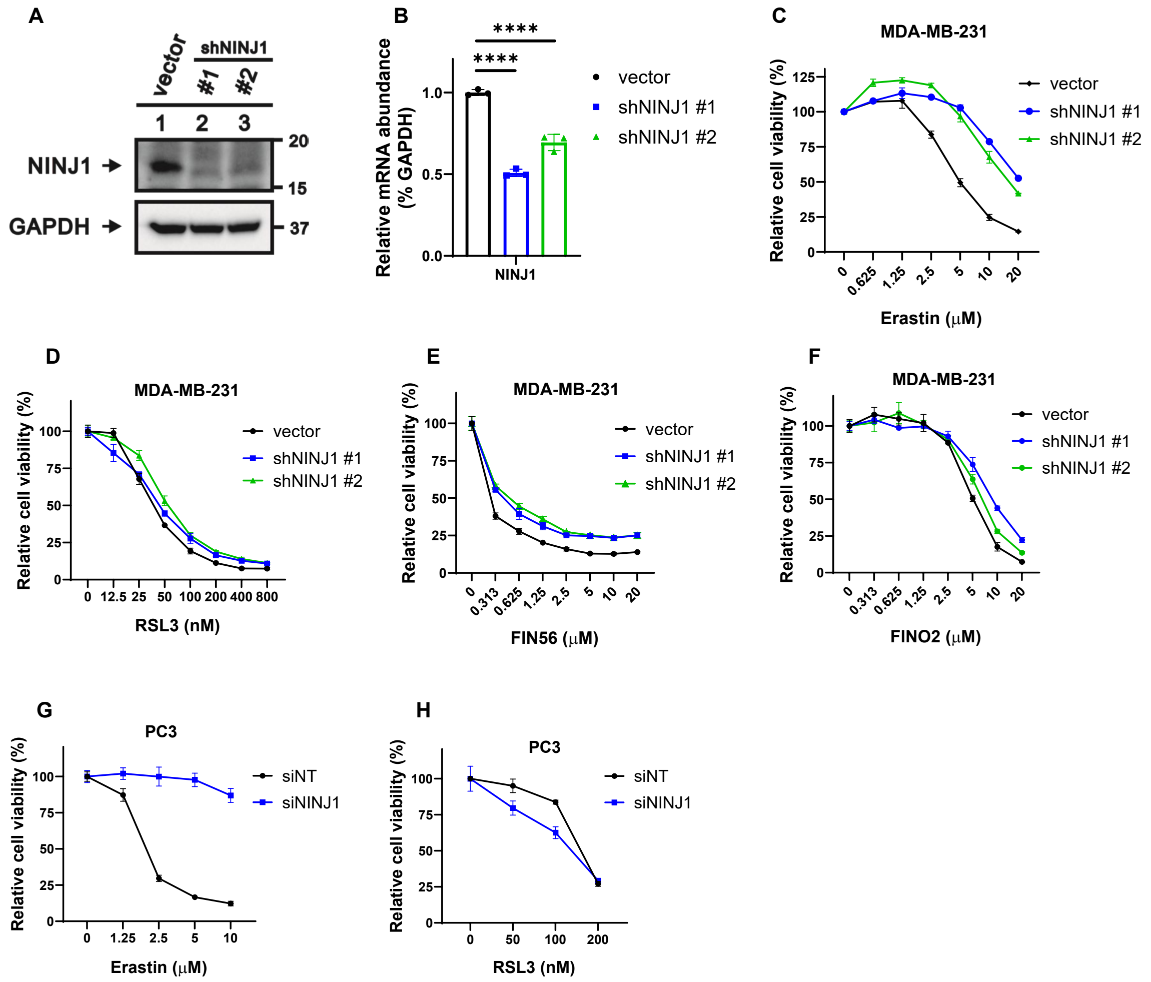


**Supplemental Figure 1.** (A-B) Two NINJ1 shRNA significantly reduced the expression of NINJ1 in HT-1080 cells verified by (A) Western blot and (B) Quantitative real-time PCR. (C-F) Cell viability of control and NINJ1-knockdown MDA-MB-231 cells were determined by CellTiter-Glo assay after cells were treated with indicated doses of (C) Erastin (26 hours), (D) RSL3 (21 hours), (E) FIN56 (21 hours), and (F) FINO2 (26 hours). (G-H) Cell viability of siNT and siNINJ1 PC3 cells was determined by CellTiter-Glo assay after cells were treated with indicated doses of (G) Erastin (28 hours) and (H) RSL3 (28 hours).

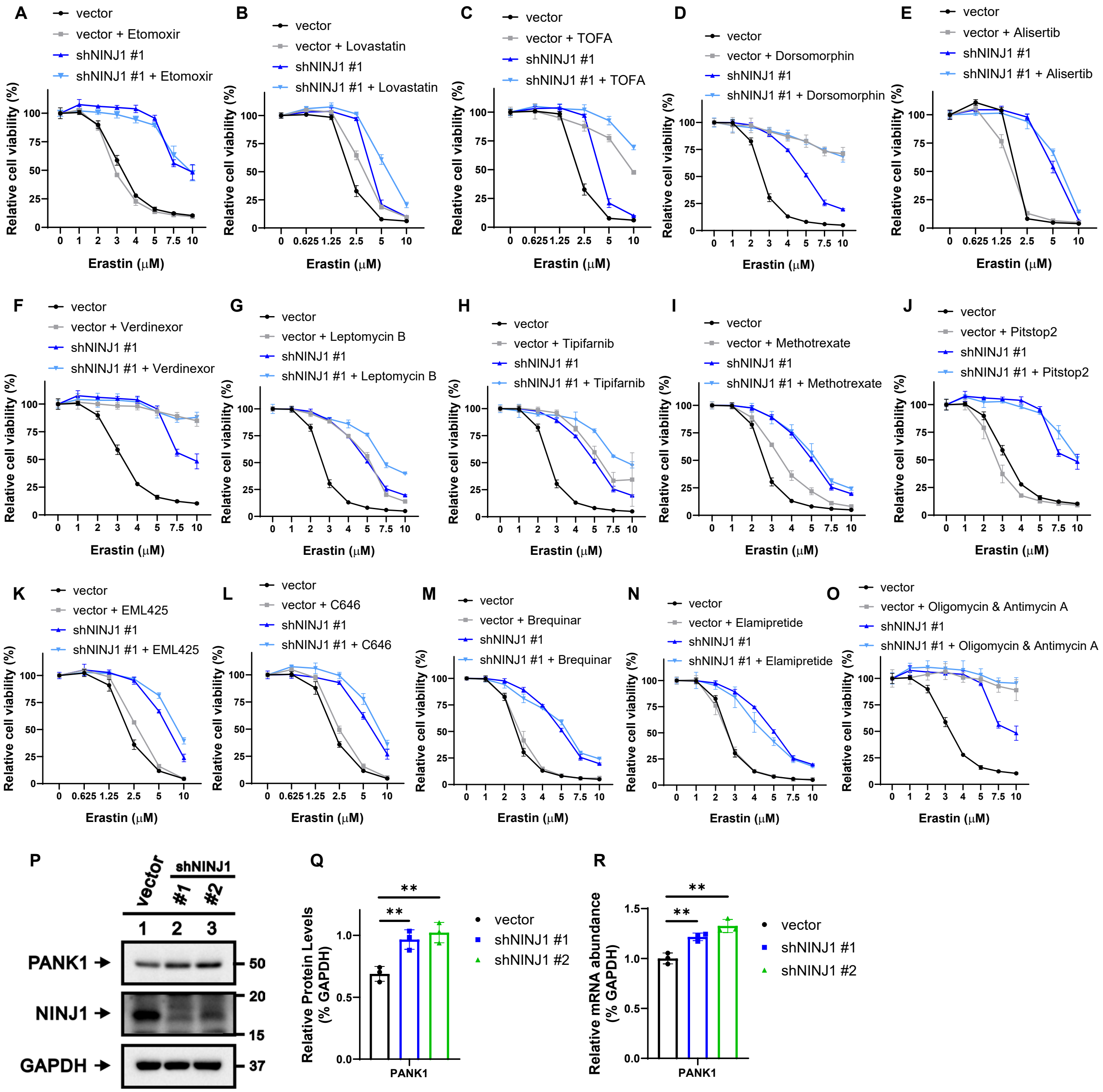


**Supplemental Figure 2.** (A-O) Cell viability of control and NINJ1-knockdown HT-1080 cells were determined by CellTiter-Glo assay after cells were treated with indicated doses of Erastin combined with/without (A) β-oxidation inhibitor (Etomoxir 5 μM, 18 hours), (B) HMGCR inhibitor (Lovastatin 10 μM, 22 hours), (C) ACC inhibitor (TOFA 25 μM, 22 hours), (D) AMPK inhibitor (Dorsomorphin 10 μM, 18 hours), (E) Aurora kinase A inhibitor (Alisertib 10 μM, 21 hours), (F-G) XPO1/CRM1 inhibitors (F Verdinexor, 18 hours & G leptomycin B 25 μg/ml, 18 hours), (H) farnesyltransferase inhibitor (Tipifarnib 10 μM, 18 hours), (I) dihydrofolate reductase inhibitor (Methotrexate 2 μM, 18 hours), (J) clathrin-mediated endocytosis inhibitor (Pitstop2 25 μM, 18 hours), (K-L) p300/CBP Inhibitor (K EML425 5 μM, 23 hours & L C646 5 μM, 23 hours), (M) DHODH inhibitors (Brequinar 500 μM, 18 hours), (N) mitochondrial-targeted peptide (Elamipretide 20 μM, 18 hours), and (O) mitophagy induction (the combination treatment of Oligomycin 10 μM and Antimycin A 10 μM, 18 hours). (A, F, J, O) were performed at the same time. (B, C) were performed at the same time. (D, G-I, M, N) were performed at the same time. Each group has the same control and NINJ1-knockdown HT-1080 cells following the treatment with indicated doses of Erastin. (P-R) NINJ1 knockdown increased PANK1 expressions verified by (P-Q) Western blot and (R) Quantitative real-time PCR.

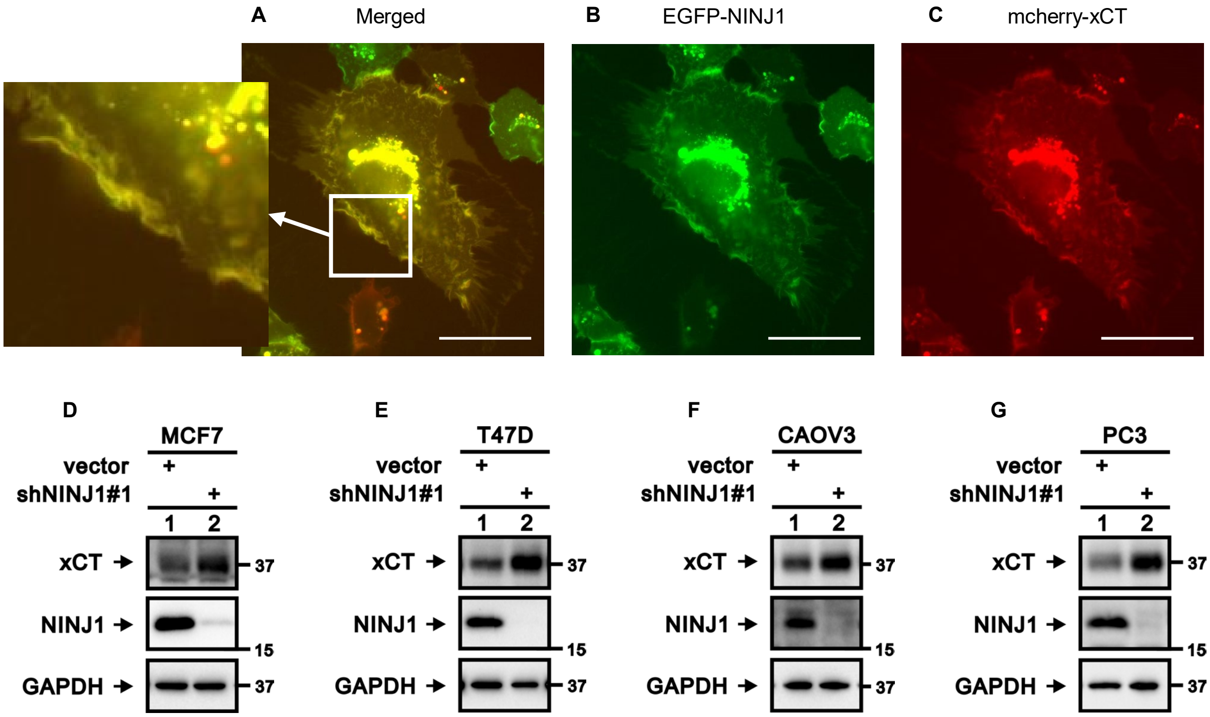


**Supplemental Figure 3.** (A-C) EGFP-NINJ1 and mCherry-xCT were overexpressed in HT-1080 cells. Fluorescent images of (A) merged, (B) EGFP-NINJ1 alone, and (C) mcherry-xCT alone are presented. Scale bar: 50 µm. (D-G) xCT expression was increased following NINJ1 knockdown in (D) MCF7, (E) T47D, (F) CAOV3, and (G) PC3 cells verified by Western blot.

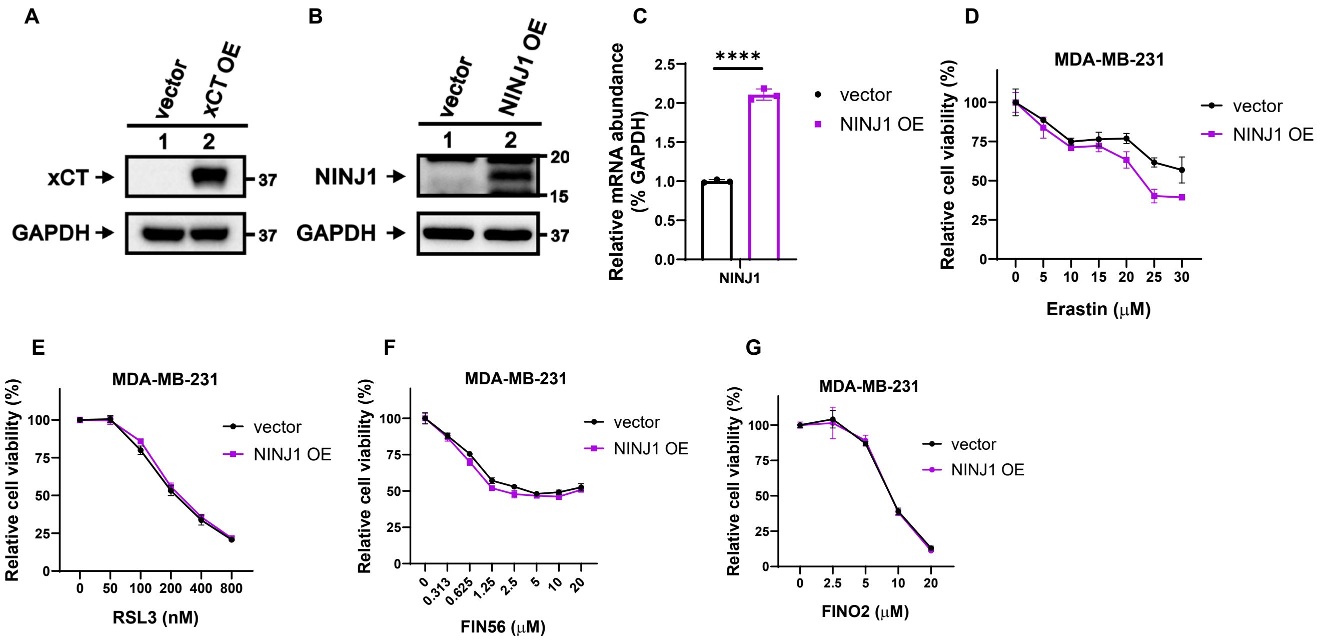


**Supplemental Figure 4.** (A) xCT overexpression efficacy in HT-1080 cells was verified by Western blot. (B-C) NINJ1 overexpression efficacy in HT-1080 cells was verified by (B) Western blot and (C) Quantitative real-time PCR. (D-G) Cell viability of control and NINJ1-overexpression MDA-MB-231 cells were determined by CellTiter-Glo assay after cells were treated with indicating doses of (D) Erastin (57 hours), (E) RSL3 (26 hours), (F) FIN56 (26 hours), and (G) FINO2 (32 hours).
